# Single-cell analysis uncovers mechanisms of plasticity in leukemia initiating cells

**DOI:** 10.1101/2020.04.29.066332

**Authors:** Vivian Morris, William Marion, Travis Hughes, Patricia Sousa, Prerana Sensharma, Yana Pikman, Marian Harris, Alex K. Shalek, Trista E. North, George Q. Daley, Edroaldo Lummertz da Rocha, R. Grant Rowe

## Abstract

Leukemia initiating cells (LICs) fuel leukemic growth and spark relapse. Previously thought to be primitive and rare, the LIC state may actually be heterogeneous and dynamic, enabling evasion of therapies. Here, we use single-cell transcriptomics to track LIC multipotency within the cellular ontogeny of *MLL*-rearranged B-lymphoblastic leukemia (*MLL*-r B-ALL). Although we identify rare transcriptionally and phenotypically primitive LICs, we also observe LICs emerging from more differentiated populations with the capability to replenish the full leukemic cellular diversity. We find that activation of MYC-driven oxidative phosphorylation controls this process of facultative state conversion in LICs.

---

Leukemia initiating cells (LICs) possess the capability for sustained, deregulated selfrenewal and the potential to fully reconstitute fulminant leukemia at relapse or upon xenotransplantation. This definition was formed based on the finding that only a specific fraction of human acute myeloid leukemia (AML) cells could engraft immunodeficient mice (*1*). Subsequent studies advanced a model wherein leukemia cells are hierarchically organized with rare, primitive, quiescent LICs at the apex that appropriate hallmarks of normal hematopoietic stem and progenitor cells (HSPCs) including selfrenewal and differentiation (*2, 3*). Based on this model, LICs have been defined in several forms of leukemia (*4–9*).

Recent advances have opened paradigms of LIC biology to revision. Use of improved immunodeficient mouse strains that heighten the sensitivity of xenotransplantation has revealed that engraftable LICs can be heterogeneous (*10–12*). This concept could explain the limited clinical success of interventions targeting LICs as has been suggested in solid tumors where cancer stem cells show phenotypic plasticity (*3, 13*). Although such findings have led to reassessment of classical LIC models, it remains generally accepted that leukemias are comprised of heterogeneous cells with variable xenotransplantation capacities. Since the engraftability of individual leukemias – a readout of LIC content - is of prognostic importance, improved understanding of LIC biology is needed (*14, 15*).

B-ALL with rearrangement of the *MLL* locus (*MLL*-r) constitutes about 80% of BALL of infancy and also occurs in older children and adults. *MLL*-r B-ALL behaves aggressively, often presenting with corticosteroid resistance and central nervous system infiltration, and has poor long-term outcomes (*16, 17*). This unfavorable clinical behavior is associated with distinctive underlying biology including coexpression of markers of myeloid differentiation and the ability to undergo a B-lymphoid-to-myeloid lineage switch (*16, 18, 19*). These attributes suggest that *MLL*-r B-ALL LICs uniquely possess primitive HSPC-like multipotency programs and multilineage differentiation potential that could be tracked to purify and study LIC biology.

Here, we used single cell RNA sequencing (scRNA-seq) combined with xenotransplantation to define the cellular diversity of *MLL*-r B-ALL. Although LICs are enriched in phenotypically and transcriptionally primitive fractions, they can facultatively emerge from more differentiated blast populations to reconstitute the full cellular diversity of *MLL*-r B-ALL, a process regulated by MYC signaling and mitochondrial oxidative phosphorylation. Our findings define new mechanisms of LIC plasticity with anticipated therapeutic relevance.

## RESULTS

### Single cell RNA sequencing identifies candidate LICs

The coexpression of myeloid markers and tendency to switch to the myeloid lineage at relapse suggests that *MLL*-r B-ALL LICs possess B-lymphoid/myeloid multipotency (*16, 19*). Therefore, we aimed to elicit multipotency at the single-cell level in *MLL*-r B-ALL. To this end, we employed patient derived xenografts (PDXs) with defined somatic mutations (Table S1)(*20, 21*). These *MLL*-r B-ALL PDXs infiltrate the liver, spleen, lymph nodes, and central nervous system of unconditioned NOD.Cg-*Prkdc^scid^Il2rg^tm1Wjl^* (NSG) recipient mice (Fig. 1A). At baseline, *MLL*-r B-ALL cells are nearly uniformly CD19^+^ with occasional CD33^+^ cells (Fig. 1B). To test for latent multipotency, we used MS5 stromal cells that can induce B-lymphoid and myeloid differentiation in human HSPCs (*22, 23*). After 4 weeks of culture on MS5, *MLL*-r B-ALL cells differentiated to CD33^+^ primitive myeloid cells (Figure S1A). By performing single cell assays, we found that *MLL*-r B-ALL cells generated clonal outgrowths containing both B-lymphoid and myeloid differentiation (Fig. S1, B-C).

**Figure. 1.**
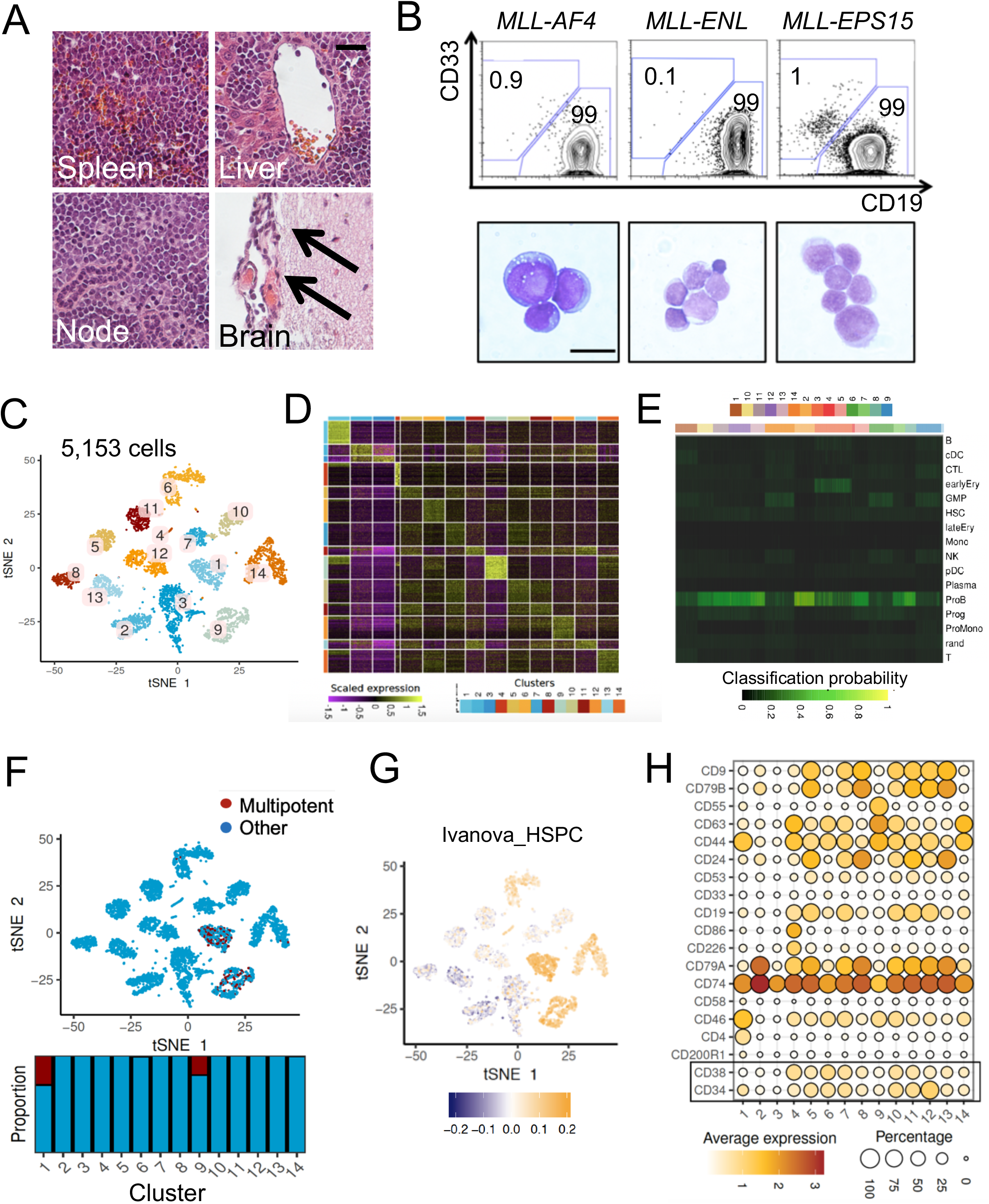
Single cell RNA sequencing identifies candidate LICs. (A) The indicated tissues were isolated from NSG mice engrafted with human *MLL*-r BALL PDXs and analyzed by light microscopy (scale = 50 μm). (B) (Upper panels) *MLL*-r B-ALL cells were procured from leukemic NSG bone marrow or spleen and analyzed by flow cytometry (gated on human CD45+). (Lower panels) *MLL*-r B-ALL cells were examined by light microscopy (scale = 10 μm; left; leukemia 1; middle leukemia 2; right; leukemia 3). (C) Human CD45+ *MLL*-r B-ALL cells were isolated from patient peripheral blood and analyzed by scRNAseq (results shown are for leukemia-1*/MLL-AF4*). t-distributed stochastic neighbor embedding (t-SNE) was used to visualize populations. (D) Differentially expressed transcripts in each cluster are shown on a heatmap. (E) SingleCellNet was used to classify individual cells relative to normal HSPC and differentiated benchmarks. (F) The StemID algorithm was used to annotate each single cell with a multipotency label, with results overlaid on the t-SNE plot and proportion of predicted multipotent cells in each cluster shown. (G) t-SNE plot showing enrichment of a validated HSC signature. (H) The expression of the indicated cell surface markers in each population is shown.

To better understand the primitive multipotent programs in *MLL*-r B-ALL, we performed scRNA-seq (*24, 25*). We obtained 5,153 viable, human CD45^+^ cells from leukemia 1 (*MLL-AF4* patient peripheral blood, inDrop platform) and 6,230 cells from leukemia 2 (*MLL-ENL* early passage PDX, SeqWell platform), after performing quality controls (see Methods). Presenting leukemia 1 (*MLL-AF4*) as an example, we visualized 14 transcriptional groups with unique signatures using t-stochastic neighbor embedding (t-SNE; Fig. 1C-D). We used the SingleCellNet algorithm to compare each subpopulation to normal benchmarks and found that most cells classified as pro-B cells, indicating that the observed transcriptional heterogeneity is independent of the global differentiation state (Fig. 1E)(*26, 27*). We next used the StemID algorithm to assign a multipotency label to each cell, finding that clusters 1 and 9 were most highly enriched in putative multipotent cells (Fig. 1F)(*28*). Consistent with this prediction, we validated enrichment of primitive HSPC signatures in these clusters (Fig. 1G)(*29*). We observed similar results in leukemia 2 (Figure S1, D-G).

To further assess the differentiation state of these candidate LSC-enriched clusters we analyzed the expression of cell surface markers. We found that the primitive HSPC marker CD34 was expressed in the candidate LIC cluster 1 along with low expression of CD38 and the committed B-cell markers CD19, CD79A, and CD79B. This suggested a relatively primitive differentiation state typical of normal human HSCs and multipotent progenitors (MPPs) in the candidate LIC cluster 1 (Fig. 1H, Figure S1, H)(*30*). Given this finding and considering the primitive multipotency programs of *MLL*-r B-ALL, we analyzed expression of CD34 and CD38 as well as the markers CD90 and CD45RA by flow cytometry. These markers resolve multipotent HSPCs - HSCs, MPPs, and multilymphoid progenitors (MLPs) - from more lineage restricted myeloid progenitors such as granulocyte/monocyte progenitors (GMPs)(*30*). We found that *MLL*-r B-ALL contained three distinguishable populations that we named based on their corresponding normal HSPC immunophenotypes: rare CD34^+^ CD38^-^ CD90^-^ CD45RA^+^ leukemic MLPs (L-MLPs), more abundant CD34^+^ CD38^+^ CD90^-^ CD45RA^+^ leukemic GMPs (L-GMPs), and the remaining CD34^-^ cells (Fig. 2A). Together, these results suggest that *MLL*-r BALL cells are heterogenous with LICs possessing a primitive phenotype and multipotency programs.

**Figure 2.**
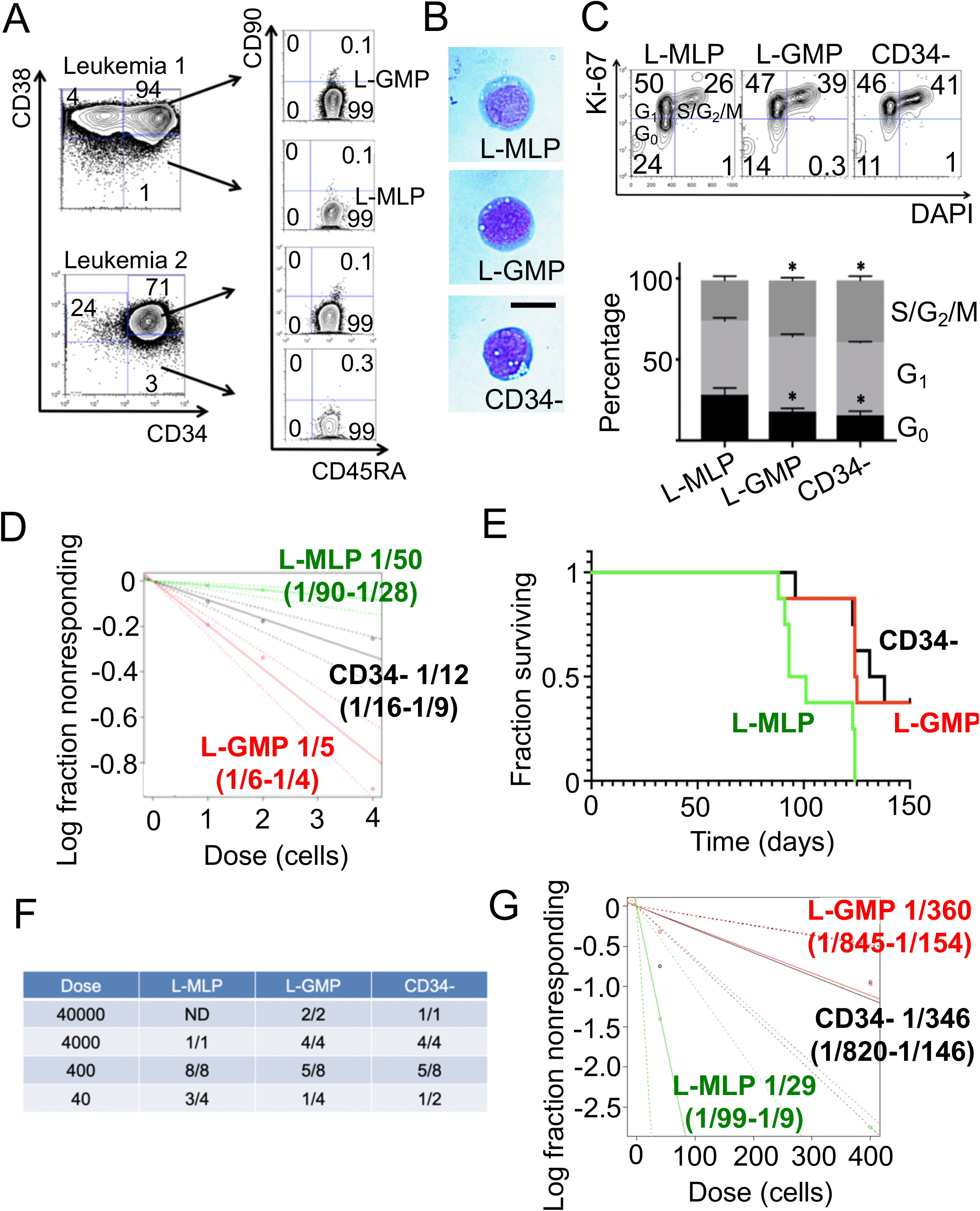
*MLL*-r B-ALL shows functional cellular heterogeneity. (A) Representative flow cytometry distributions of cells based on the indicated markers. (B) The indicated populations were sorted from leukemic marrow (leukemia *1/MLL-AF4*) and morphology examined (scale = 10 μm). (C) The indicated populations were isolated by FACS and cell cycle state analyzed by flow cytometry following staining for Ki67 and DNA content with DAPI. Cell cycle distribution of each population was quantified (results aggregated over two independent experiments, n = 5 xenografted mice tested; * p < 0.05 compared to L-MLPs by unpaired student’s t-test). (D) The indicated populations were sorted onto MS5 stromal layers at various doses. After 4 weeks, outgrowths were tabulated, with estimated progenitor cell frequency presented. Results are aggregated over two independent experiments (for L-MLP versus L-GMP X^2^ = 64.3, p = 1 x 10^-15^; for L-MLP versus CD34^-^ X^2^ = 14.9, p = 0.0001; for L-GMP versus CD34^-^ X^2^ = 18.1, p = 0.00002). (E) 400 cells of the indicated fractions were xenotransplanted into unconditioned NSG recipients, and the incidence of leukemia monitored over time (results aggregated over two independent transplantation experiments; p = 0.008 L-MLP versus L-GMP, 0.005 L-MLP versus CD34^-^, and 0.9 L-GMP versus CD34^-^ compared by log rank test). (F-G) LIC content of each population was quantified by in vivo LDA with incidence at each dose shown (see Table S2).

### *MLL*-r B-ALL shows functional cellular heterogeneity

To determine if *MLL*-r B-ALL LICs are enriched in any of these three populations, we used fluorescence activated cell sorting (FACS) to purify each for analysis (Fig. 2B, Fig. S1, I). Since LICs are typically quiescent, we analyzed the cell cycle status of each population (*31–33*). The most primitive L-MLP fraction contained the highest proportion of cells in G_0_ phase (Fig. 2C, Fig. S2, A). Next, we used MS5 assays to quantify leukemic progenitors, finding that L-GMPs possessed the highest frequency of shortterm clonogenic cells (Fig. 2D). As the gold standard to detect LICs, we next used limiting dilution analysis (LDA) xenotransplantation (*34*). Using the terminal leukemia as our endpoint, we first observed that unfractionated human *MLL*-r B-ALL possessed a remarkably high frequency of LICs (1/426 cells (1/1417 – 1/128 95% confidence interval)). Using FACS to fractionate *MLL*-r B-ALL PDX cells, we found that L-MLPs caused leukemia with the shortest latency and contained the highest LIC content in LDA, an effect we corroborated with primary patient cells (Fig. 2E-G, Fig. S2, C-H, Fig. S3, A-B, Table S2). In line with this finding, using RNA sequencing (RNA-seq) of FACS-purified populations, we found that the L-MLP fraction bore the strongest primitive HSPC signature, paralleling our scRNA-seq data (Fig. S3, C-D).

Since quiescent LICs may be relatively resistant to chemotherapy, we exposed *MLL*-r B-ALL cells to prednisolone for five days followed by a washout period to monitor leukemic regeneration, mimicking the corticosteroid prophase used in chemotherapy regimens (*16*). While most cells died upon prednisolone exposure, we found that L-MLPs were most enriched in the surviving fraction (Fig. S3, E-G). Over three weeks of culture, we observed that surviving cells underwent a myeloid lineage switch (Fig. S3, H-J). Together, these results demonstrate that *MLL*-r B-ALL cells are functionally heterogeneous and appear to be hierarchically organized, with primitive, corticosteroid-resistant, phenotypically plastic, LIC-enriched L-MLPs at the apex.

### Plasticity of *MLL*-r B-ALL LICs

Although the primitive L-MLP fraction contained abundant LICs, we observed that terminal leukemia did occur in recipients of the more differentiated L-GMP and CD34^-^ fractions, albeit at lower efficiency (Fig. 2E-G). When we compared the content of terminal leukemia derived from each fraction, we found that the full diversity emerged from each source (Fig. 3A). Moreover, by secondary xenotransplantation, we found that leukemia derived from each source contained serially transplantable, self-renewing LICs (Fig. 3B). To directly observe this plasticity during leukemic reconstitution in vivo, we transplanted purified CD34^-^ cells into NSG mice and isolated bone marrow at defined time points post-transplant but prior to the expected onset of terminal leukemia. We found that CD34-cells gradually replenished the CD34+ fraction over time (Fig. 3C). To exclude an artifactual effect of the PDX model, we used primary patient blasts and observed similar LIC plasticity in vivo and down to the single-cell level in vitro, where we also observed latent myeloid potential in single-cell clones from all phenotypic populations (Fig. S4, A-D).

**Figure 3.**
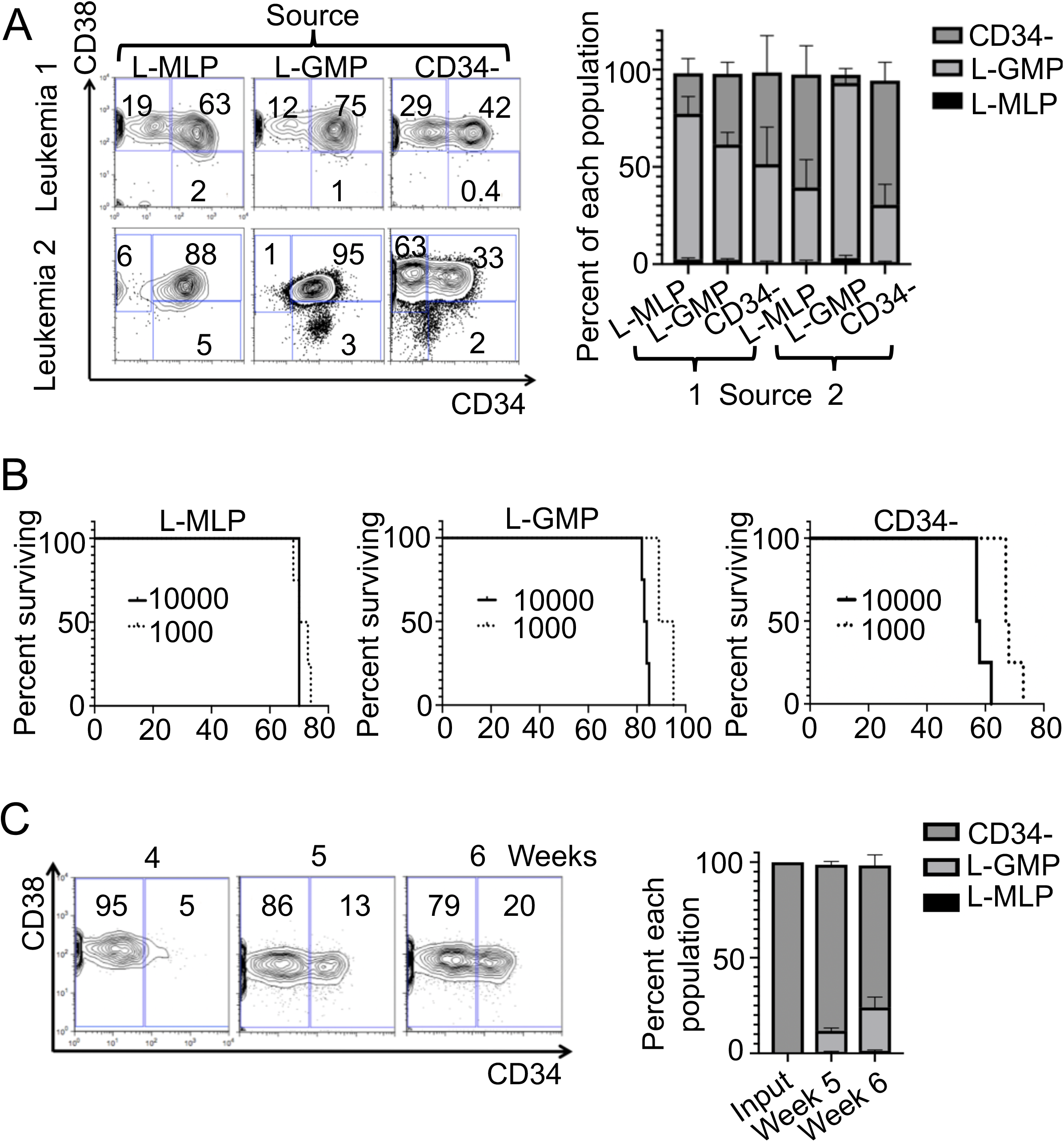
Plasticity of *MLL*-r B-ALL LICs. (A) Flow cytometry profiles of terminal leukemias derived from the indicated transplanted cell populations gated on viable and human CD45+ cells within bone marrow. Proportions of each cell type in leukemias derived from the indicated transplanted cell populations are presented (p = NS comparing each outcome population except in leukemia 1 where p < 0.05 comparing the L-GMP and CD34^-^ fractions of leukemias derived from L-GMP to both L-MLP and CD34^-^ derived leukemia). (B) Terminal leukemias derived from the indicated cell sources of leukemia *1/MLL-AF4* in primary transplantation were transplanted into secondary recipients at the indicated doses, and the onset of terminal leukemia was monitored in the secondary recipients. (C) CD34^-^ cells were purified from leukemia 1 by FACS and transplanted into recipient mice. Reconstitution of CD34^+^ populations was monitored by flow cytometry (gated on viable human CD45^+^ cells) and quantified over time.

### LIC plasticity is driven by *MYC* and oxidative metabolism

To investigate mechanisms of LIC plasticity, we performed RNA-seq on relatively LIC-deplete CD34^-^ cells either transplanted alone and actively repopulating CD34^+^ populations or growing at steady state in vivo with L-GMPs and L-MLPs (Figure 4A). Despite their identical immunophenotype, regenerating CD34^-^ cells (CD34^-^r) and steadystate CD34^-^ cells bore divergent transcriptional profiles (Figure 4B). We found that CD34^-^r cells bore signatures of oxidative phosphorylation and MYC target gene expression (Figure 4C). At steady state, these signatures are enriched in multipotent scRNA-seq clusters, suggesting that CD34^-^r cells are activating primitive programs (Figure 4D). Using available MYC ChIP-seq data, we found that MYC bound loci involved in mitochondrial function and oxidative metabolism (encodeproject.org, Figure 4E).

**Figure 4.**
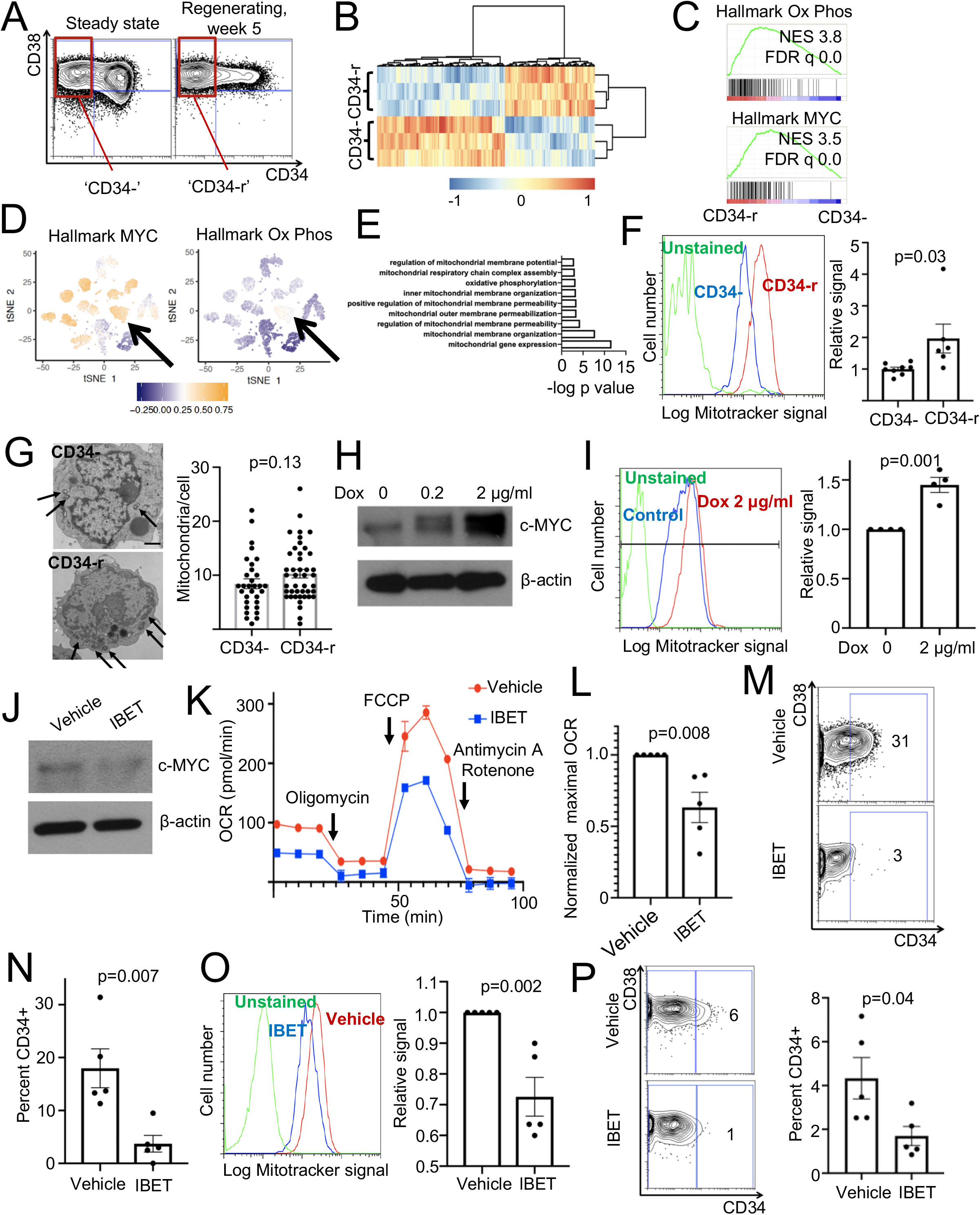
LIC plasticity is driven by *MYC* and oxidative metabolism. (A) Schematic showing the input populations sorted for RNA-seq at steady state (left) and from regenerating CD34^-^ cells (right). (B) RNA-seq was performed on the indicated populations. Representative heatmap of differentially expressed gene is shown across three replicates. (C) GSEA was used to identify differentially enriched signatures in CD34^-^r versus CD34^-^ cells. (D) The indicated expression signatures were analyzed in the context of the corresponding scRNA-seq data. (E) MYC ChIP-seq data from the ENCODE project were analyzed and significantly enriched peaks queried using Gene Ontology analysis, with significant terms presented. (F) Active mitochondria were quantified in the CD34^-^ fraction of bulk leukemia or CD34^-^r cells using Mitotracker green, with representative flow cytometry results presented compared to background of unstained cells (n = 8 CD34^-^ and 6 CD34^-^r biologic replicates over two independent experiments). (G) CD34^-^ or CD34^-^r cells were processed for transmission electron microscopy (TEM), with representative images presented (scale = 1 μm). Number of mitochondria per cell were quantified over two experiments. (H) MV4;11 cells were transduced with a doxycycline inducible MYC vector and treated with the indicated doses of doxycycline for 24 hours, at which time MYC was measured by Western blotting. (I) MYC expression was induced in MV4;11 cells for 4 days, at which time mitochondrial activity was measured by flow cytometry using Mitotracker (n = 4 independent experiments). (J) MV4;11 cells were treated for 48 hours with 50 n*M* IBET-151 at which time MYC was measured by Western blot. (K-L) MV4;11 cells were treated with 50 n*M* IBET-151 or vehicle control for 48 hours, at which time they were analyzed by the Mito Stress Assay. Plot is representative of five independent experiments analyzed in (L). (M-O). FACS-sorted CD34^-^ cells were cultured on MS5 stroma for 14 days under the indicated conditions, at which time either the CD34^+^ content (M) or Mitotracker green signal (N) of the cultures was analyzed by flow cytometry (n = 5 biologic replicates). (P) CD34-r cells were engrafted into NSG mice for three weeks, at which time a two week treatment course with IBET-151 (5 mg/kg IP daily Monday-Friday for two weeks), after which the human CD45+ content of the bone marrow was analyzed by flow cytometry with the indicated markers (n = 5 mice per group). All results presented as mean ± SEM and compared by student’s t-test, with p-values for the indicated comparisons shown.

Since MYC activity drives mitochondrial oxidative metabolism and is implicated in the pathobiology of *MLL*-r leukemias, and since LICs rely on oxidative phosphorylation, we investigated the role of this pathway in LIC plasticity (*35–37*). By using the Mitotracker assay, we confirmed that CD34^-^r cells contained higher mitochondrial activity compared to CD34^-^ cells, although overall mitochondrial mass was similar (Figure 4F-G). Next, we generated MV4;11 *MLL*-r biphenotypic B-ALL/myeloid cells bearing a conditional, doxycycline-inducible MYC expression cassette (Figure 4H)(*38*). Activation of MYC expression was sufficient to increase mitochondrial metabolism (Figure 4I). MYC expression can be inhibited by the bromodomain inhibitor IBET-151 (*39*). Therefore, we treated MV4;11 cells with IBET-151 and confirmed that this reduced MYC protein (Figure 4J). By using the Seahorse Mito Stress Assay, we found that IBET-151 diminished mitochondrial oxidative metabolism (Figure 4K-L). Next, we treated FACS-sorted CD34^-^r cells in culture on MS5 and actively undergoing regeneration with IBET-151 and monitored state plasticity. We found that IBET-151 inhibited conversion to the CD34^+^ state in culture and diminished active mitochondrial content (Figure 4M-O). Treatment of NSG mice transplanted with CD34-r cells impaired acquisition of a CD34+ phenotype (Figure 4P). These results demonstrate that MYC regulates the metabolic state of *MLL*-r B-ALL cells as a mechanism of cell state interconversion.

## DISCUSSION

Since the initial supposition of malignant stem cells, models of leukemic ontogeny have been predicated upon that of normal hematopoiesis (*2–4*). However, the classical model holding that LICs are uniformly primitive akin to normal HSCs have gradually been revised (*3*). At first glance, our data would appear to be consistent with this classical model in that *MLL*-r B-ALL cells are stratified from phenotypically and transcriptionally primitive to most differentiated. However, in B-ALL, LICs are distributed across multiple phenotypic populations, although their relative enrichment in each apparent differentiation state had not been previously quantified (*1, 8, 9, 40, 41*). Compared to earlier studies, use of more immunodeficient mouse models with heightened sensitivity for human engraftment has uncovered heterogeneity in AML LICs as well (*11, 42*). Although across many forms of leukemia, LIC ontogeny does not seem to follow the strict stratification and unidirectional differentiation of normal HSPCs, the idea that engraftable LIC content correlates with leukemia prognosis provides an impetus for investigation of the molecular determinants of their state (*14, 15*). Notably, we find that in *MLL*-r B-ALL, functional LICs are quite common, being present in approximately 1/500 cells, which is far more frequent than has been reported in limiting dilution of adult AML (*11*).

Our data reinforce the notion that LIC identity can be fluid. Use of NSG mice led to the first evidence that AML LICs within CD34^-^ fractions could reconstitute the full cellular diversity of the original AML specimen, including more primitive CD34^+^ cells, although the mechanism of this interconversion was not known (*11*). In a clinical context, in a more recent study, use of matched diagnostic and relapse AML patient specimens revealed that the frequency of engraftable LICs expanded up to 90-fold and that LICs gained heterogeneity at relapse, suggestive of plasticity under chemotherapeutic pressure (*12*). Moreover, a recent study reported that acute promyelocytic leukemia cells undergoing differentiation driven by retinoic acid could re-acquire the LIC state following chemotherapy withdrawal, effectively reversing the differentiation trajectory (*10*). Extending this concept to solid tumors, a similar effect was recently uncovered in colon cancer, where cells devoid of the stem cell marker Lgr5 can establish metastases and give rise to Lgr5^+^ cancer stem cells (*43*). These findings parallel the known phenomenon that in healthy tissues, adult stem cell identity can be plastic: professional stem cells maintain tissue integrity at steady state, but under stress, facultative stem cells can be recruited from apparently lineage-committed populations (*44, 45*).

Our efforts to uncover mechanisms of LIC plasticity in *MLL*-r B-ALL implicated MYC signaling and oxidative phosphorylation. Oncogenic MYC activity is a downstream effector of transforming MLL translocations and mitochondrial turnover is important for LIC homeostasis (*39, 46*). Interestingly, MYC can drive mitochondrial oxidative metabolism and proliferation of stem cells in breast cancer, consistent with our findings (*35*). Recruitment of oxidative metabolism seems to also play a role in relapse in B-ALL (*47*). We impaired MYC activity through inhibition of BET bromodomain proteins - an intervention that slows progression of *MLL*-driven leukemia - finding that this inhibitor blocked the conversion of CD34^-^ to CD34^+^ cells (*39, 48, 49*). Together with prior studies, our data support a model wherein MLL-containing translocations cooperate with bromodomain proteins to engage MYC and activate oxidative phosphorylation, driving LIC proliferation and repletion of all leukemic populations. Our findings can be connected to the clinical behavior of *MLL*-r B-ALL and may have therapeutic implications. The dual lineage and LIC plasticity of *MLL*-r B-ALL provide two distinct mechanisms of chemotherapy evasion. Together with high LIC content, this promiscuity in cell state likely contributes to the poor outcomes of *MLL*-r B-ALL. Further understanding of the molecular basis of the enhanced cellular plasticity in *MLL*-r B-ALL relative to normal hematopoiesis could lead to valuable therapies.

Together, our findings reinforce the notion that LICs are plastic and adaptable, providing possible explanations as to why therapies targeting LICs have yet to prove widespread efficacy despite being the object of intense investigation for over two decades (*3, 50*). Although aspects of LICs in *MLL*-r B-ALL might prove to be disease-specific, placement of our results in the broader context of the prevailing knowledge of LICs illustrates the ongoing revision of classical LIC paradigms. Further incremental innovation in immunocompromised mouse models and single-cell-level readouts could continue to augment understanding of LIC biology and define an as yet evasive unifying LIC model.

## Supporting information

Table S2 part 1

Table S2 part 2

## ACKNOWLEDGEMENTS

The authors thank the Single Cell Core at Harvard Medical School, Boston, MA for performing the single-cell RNA-Seq sample preparation. We thank the Molecular Biology Core Facility at the Dana-Farber Cancer Institute for sequencing studies. We thank William Oldham at the Seahorse Core at Brigham and Women’s Hospital. We thank the Flow Cytometry Core at Boston Children’s Hospital. This work was supported by the National Institute of Diabetes and Digestive and Kidney Diseases (K08 DK114527-01 to R.G.R.) as well as grants from the St. Baldrick’s Foundation, Pablove Foundation, and Pedals for Pediatrics (to R.G.R.); and the National Heart, Lung, Blood Institute (U01 HL134812 to G.Q.D.) and the Leukemia and Lymphoma Society of America (to G.Q.D.). We thank Scott Armstrong for helpful discussions.

## AUTHOR CONTRIBUTIONS

Conceptualization, R.G.R and E.L.d.R.; Methodology, R.G.R and E.L.d.R.; Software, E.L.d.R.; Formal analysis, R.G.R and E.L.d.R.; Investigation, V.M., E.L.d.R., W.M., T.H., P.S., P.S., and R.G.R.; Resources, Y.P., M.H., and A.K.S.; Data curation, T.H., R.G.R and E.L.d.R.; Writing – original draft, R.G.R. and E.L.d.R.; Writing – revising and editing, A.K.S., T.E.N., and R.G.R.; Supervision, T.E.N., G.Q.D., and R.G.R.; Project administration, T.E.N., G.Q.D., and R.G.R.; Funding acquisition, T.E.N., G.Q.D., and R.G.R.

## DECLARATION OF INTERESTS

The authors declare no competing interests.

## METHODS

### Study design

This study was initially designed to define LICs in *MLL*-r B-ALL, relying on xenotransplantation with LDA and terminal leukemia incidence by day 150 posttransplant as endpoints. This endpoint was chosen based on the approximately 100-day latency of bulk transplanted leukemia cells at a dose of 10,000 cells per transplant. We hypothesized that transplantation of LIC-enriched populations would shift latency earlier and transplantation of LIC-depleted would shift the latency further. Leukemia populations were isolated and xenotransplanted with investigators monitoring recipient mouse health blinded to the cell source. At the earliest onset of discernible morbidity suggestive of active leukemia, mice were humanely euthanized to harvest tissues and bone marrow. At the completion of the experiment, the experimental conditions were unblinded, and LIC frequency calculated by LDA analysis, or survival analyzed by logrank test, where indicated. For quantification of flow cytometry or expression data, the statistical tests used are indicated. Student’s t-tests were used and analysis was unpaired, except where otherwise indicated.

### Mice and xenotransplantation

Unconditioned NSG mice (Jackson Laboratory stock 005557) were transplanted with the indicated cell sources at the indicated doses. All transplantation was performed by tail vein injection. Mice transplanted with patient derived xenograft cells or primary patient cells were unconditioned. Mice were followed to the onset of terminal leukemia as indicated in the study design.

### Cell culture

Patient derived leukemia cells were cultured on MS5 stromal cells in the presence of 50 ng/ml recombinant human stem cell factor (SCF), 50 ng/ml recombinant human thrombopoietin (TPO), 10 ng/ml recombinant human FLT3 ligand (FLT3L), and 10 ng/ml recombinant human interleukin-7 (IL-7, all from R and D Systems)(*23*). Leukemia-1 was most amenable to in vitro culture and so was used for most culture-based experiments. For in vitro LDA experiments, 5,000 MS5 cells were plated in wells of gelatin-coated Nunc 96 well plates (Fisher Scientific) with cytokines 48 hours prior to FACS-based sorting of leukemia cells directly into the wells. Where indicated, cells were treated with 50 μg/ml prednisolone (Sigma) for the duration described.

Human MV4;11 cells were cultured in RPMI with 10% fetal calf serum supplemented with penicillin and streptomycin. Cells were treated with 0.2-2 μg/ml doxycycline hyclate (Sigma) where indicated.

### Flow cytometry and cell sorting

Human antibodies used for flow cytometry studies are listed in the resources table. Data were acquired on either BD LSR Fortessa or LSR II Instruments (BS Biosciences). Cells were sorted on a BD FACS Aria (BD Biosciences) with a 100 μm nozzle. Mitotracker Green was purchased from Thermo.

### RNA sequencing

For scRNA-Seq using the Seq-Well platform, 20,000 cells were applied to Seq-Well devices pre-loaded with mRNA capture beads as previously described(*24*). Following hybridization and reverse transcription, random second-strand synthesis was performed to generate double stranded cDNA. PCR was performed using the following primer sequence 5’ – AAGCAGTGGTATCAACGCAGAGT – 3’. Sequencing libraries were generated using the Illumina Nextera XT protocol using custom N700 sequencing indices. Libraries were sequenced using Next-Seq 75 cycle high output sequencing kits with 20 base read 1 sequence and 50 base read 2 sequence.

For inDrops-seq the cells were encapsulated in 2-3 nl droplets using a microfluidic device and the libraries were made following a previously described protocol(*25, 51*), with the following modifications in the primer sequences. RT primers on hydrogel beads- 5’CGATTGATCAACGTAATACGACTCACTATAGGGTGTCGGGTGCAG[bc1,8nt]GTCT CGTGGGCTCGGAGATGTGTATAAGAGACAG[bc2,8nt]NNNNNNTTTTTTTTTTTTTTTT TTTV- 3’

R1-N6 primer sequence (step 151 in the library prep protocol in (*51*))- 5’TCGTCGGCAGCGTCAGATGTGTATAAGAGACAGNNNNNN-3’

PCR primer sequences (steps 157 and 160 in the library prep protocol in(*51*))- 5’-AATGATACGGCGACCACCGAGATCTACACXXXXXXXXTCGTCGGCAGCGTC-3’, where XXXXXX is an index sequence for multiplexing libraries.

5’- CAAGCAGAAGACGGCATACGAGATGGGTGTCGGGTGCAG-3’

With these modifications in the primer sequences, custom sequencing primers are no longer required. Single-cell RNA-Seq was library preparation was performed by the Single Cell Core at Harvard Medical School, Boston, MA.

### Bulk RNA sequencing library prep

Cells were sorted by FACS and lysed in Trizol reagent (Thermo). RNA was isolated using RNAeasy columns (Qiagen) and low input libraries prepared in collaboration with the Molecular Biology Core at the Dana-Farber Cancer Institute.

### Data analysis

#### InDrop Single-cell RNA-sequencing

Raw sequencing reads were processed using the inDrop pipeline (https://github.com/indrops/indrops) using default parameters(*25*) The GRCh38 reference genoma was used for alignment of sequencing reads. We used scImpute to account for dropout rates in single-cell RNA-seq data and obtain an imputed count matrix that was used for all downstream analysis described(*52*). We used scImpute with the parameter ‘Kcluster = 10’. To analyze imputed single-cell inDrop data we performed quality control, dimensionality reduction, clustering and differential expression analysis using CellRouter(*53*). For this leukemia, we applied the following quality control metrics: all genes that were not detected in at least 20 cells were excluded. All cells with less than 200 genes detected were also excluded. As expression of ribosomal or mitochondrial genes is indicative of technical variation in single-cell RNA-seq data we also removed cells where the proportion of the transcript counts derived from mitochondrial genes was greater than 10%(*54*). After such quality control of the imputed count matrix, we retained 5,153 cells with a median of 15,214 genes detected per cell.

The data was then scaled and used for dimensionality reduction. We performed a principal component (PC) analysis using all genes (34,747 genes) and selected the top 20 PCs using the elbow method. These PCs were used for graph-based clustering to identify clusters of transcriptionally similar cells in our dataset. We also used the top 20 PCs to perform spectral t-stochastic neighbor embedding (t-SNE) analysis and visualize the underlying cluster structure in a space of reduced dimensionality.

#### SeqWell single-cell RNA-sequencing

Read alignment was performed as described(*55*). Briefly, for each NextSeq sequencing run, raw sequencing data was converted to FASTQ files using bcl2fastq2 that were demultiplexed by Nextera N700 indices corresponding to individual samples. Reads were first aligned to HgRC19, and individual reads were tagged according to the 12-bp barcode sequence and the 8-bp UMI contained in read 1 of each fragment. Following alignment, reads were binned and collapsed onto 12-bp cell barcodes that corresponded to individual beads using Drop-seq tools (http://mccarrolllab.com/dropseq). Barcodes were collapsed with a single-base error tolerance (Hamming distance = 1), with additional provisions for single insertions or deletions. An identical collapsing scheme (Hamming distance = 1) was then applied to UMIs to obtain quantitative counts of individual mRNA molecules. We also used scImpute to impute the raw counts matrix obtained, with the parameter of ‘Kcluster = 6’. For this leukemia, we removed all genes expressed in less than 10 cells and also removed all cells expressing less than 500 genes. Cells with transcript counts derived from mitochondrial genes larger than 10% were also removed. After QC, we retained 6,320 cells with a median of 2,588.5 genes detected per cell.

#### Normalization

Both inDrop and Seq-Well data were analyzed with CellRouter. In CellRouter, transcript counts are normalized using a global scaling normalization method that normalizes expression measurements for each cell by the total expression, multiplied by a scale factor of 10,000, and log-transformed the result.

#### StemID analysis

First, with the raw counts data obtained from the inDrop sample, we performed an initial quality control removing cells not expressing at least 200 genes or genes not expressed in at least 20 cells. This filtered, not normalized counts matrix, was used as input for scImpute with “Kcluster=10”. After data imputation, we used the imputed count matrix for StemID analysis. Briefly, we removed cell cycle genes and performed the StemID analysis setting the following parameters: mintotal=0.01, minexpr=0, minnumber=0, maxexpr=Inf, downsample=TRUE, dsn=1 in the “filterdata” function, outminc=5,outlg=2,probthr=1e-3,thr=2**-(1:40),outdistquant=.95 in the “findoutliers” function.

#### SingleCellNet analysis

We downloaded scRNA-seq data from the GEO accession number GSE116256(*27*). This study performed a random sampling of hematopoietic cells in the normal and leukemic bone marrow (BM) ecosystem. We reanalyzed five healthy BM samples published with this study and used cell types identified by the authors to train machine learning models of cell type identity of BM cells using SingleCellNet. After training, we classified each single cell in our leukemia samples as belonging to any of the classes in our training dataset.

#### Signature Scores

We downloaded gene lists from https://www.gsea-msigdb.org/gsea/msigdb. Specifically, we downloaded the following gene sets: HALLMARK_MYC_TARGETS_V1.txt, HALLMARK_OXIDATIVE_PHOSPHORYLATION.txt and IVANOVA_HEMATOPOIESIS_STEM_CELL.txt. Then, we used CellRouter to calculate signature scores for each cell and plotted the distribution of these scores.

#### Bulk RNA-sequencing

Fastq files containing single-end RNASeq reads were aligned with Tophat 2.0.12 against the UCSC hg28 reference genome using Bowtie 2.2.4 with default settings(*56, 57*). Gene level counts were obtained using the subRead featureCounts program (v1.5.1) using the parameter “--primary” and gene models from the UCSC hg28 Illumina iGenomes annotation package(*58*). Read counts were normalized using size factors as available by the DESeq2 package(*59*).

### Morphology

For morphologic analysis, leukemia cells were spun onto slides and stained with May-Grunwald and Giemsa stains (Sigma) sequentially. Transmission electron microscopy was performed at the electron microscopy core at Harvard Medical School.

### Recombinant DNA

The human *MYC* cDNA was purchased from Addgene (pDONR223_MYC_WT, a gift from Jesse Boehm and Matthew Meyerson and David Root (plasmid # 82927; http://n2t.net/addgene:82927; RRID: Addgene_82927). The *MYC* cDNA was cloned into the pCW57.1 vector (gift from David Root (Addgene plasmid # 41393; http://n2t.net/addgene:41393; RRID:Addgene_41393) using LR clonase (Thermo). The purified plasmid was used to generate lentivirus in HEK-293T cells, which was used to transduce MV4;11 cells. A stable transduced polyclonal line was selected with 0.5 μg/ml puromycin (Thermo). Gene expression was confirmed by Western blotting following doxycycline exposure

### Seahorse Assay

The Seahorse assays were performed using the Agilent Seahorse XF Cell Mito Stress Test kit using injections of 1 μ*M* oligomycin, 1 μ*M* carbonyl cyanide-4 (trifluoromethoxy) phenylhydrazone (FCCP) and 0.5 μ*M* each of antimycin A and rotenone at the intervals indicated. The instrument is located at the Seahorse Core at Brigham and Women’s Hospital.

---

**Figure S1.**
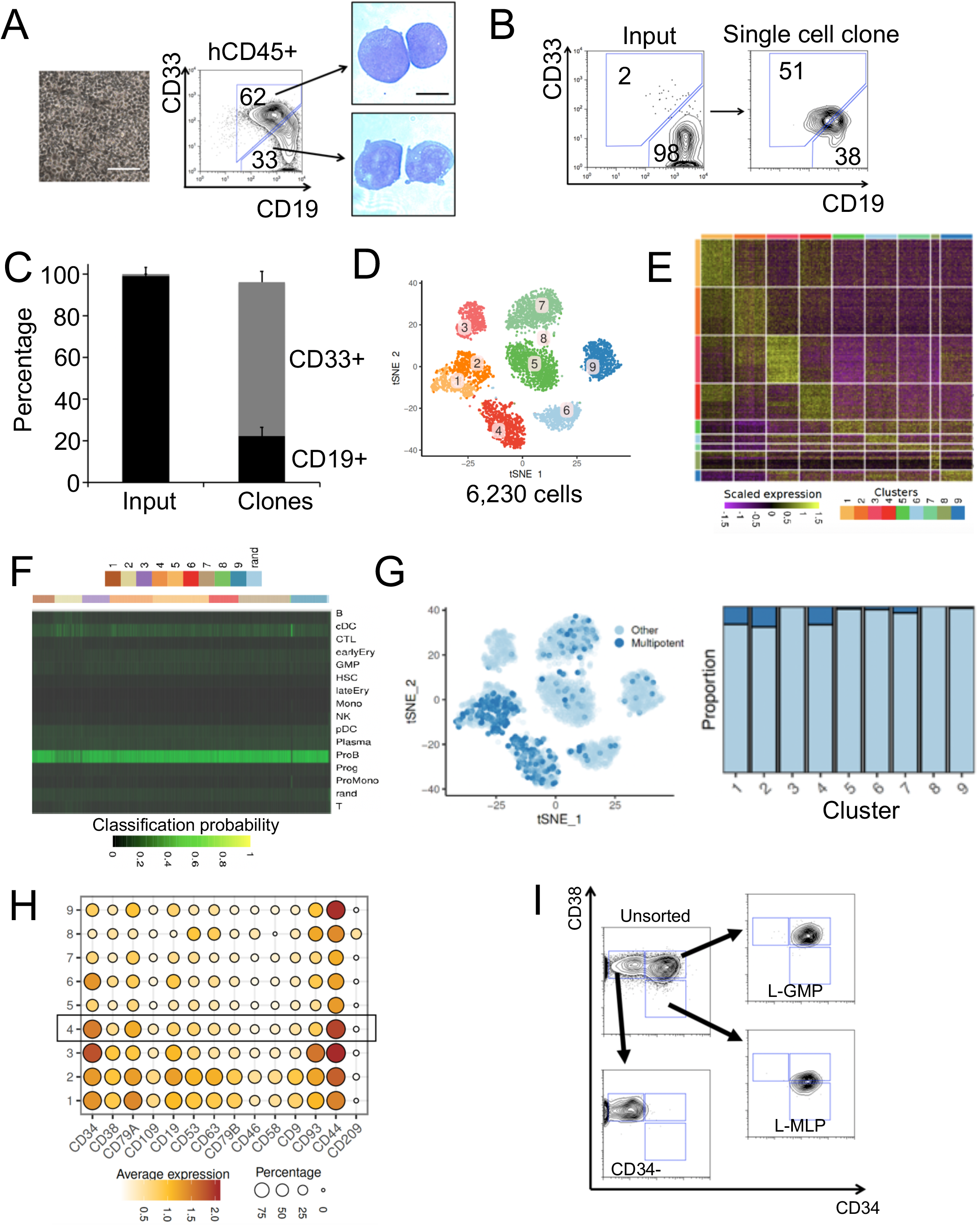

**Figure S2.**
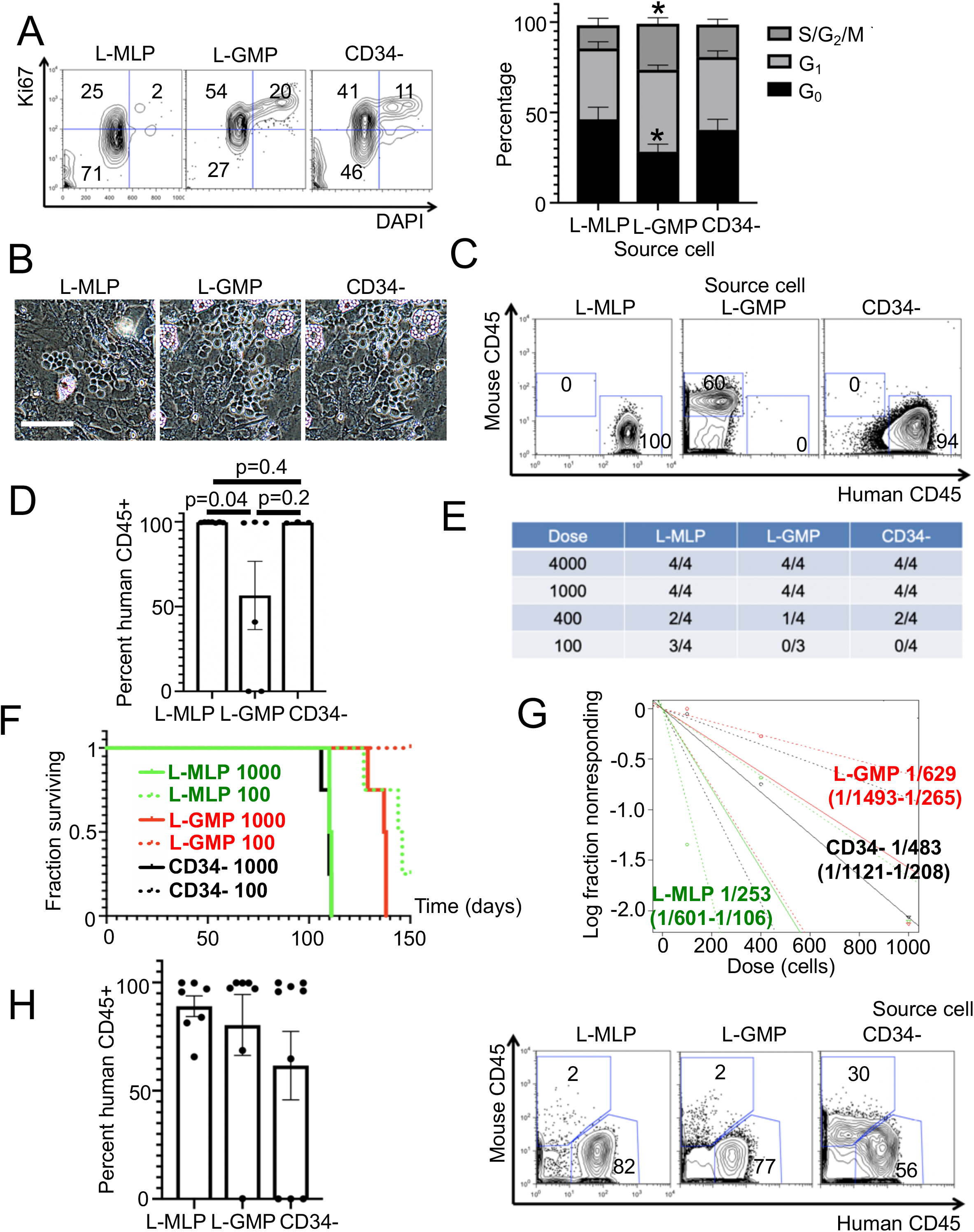

**Figure S3.**
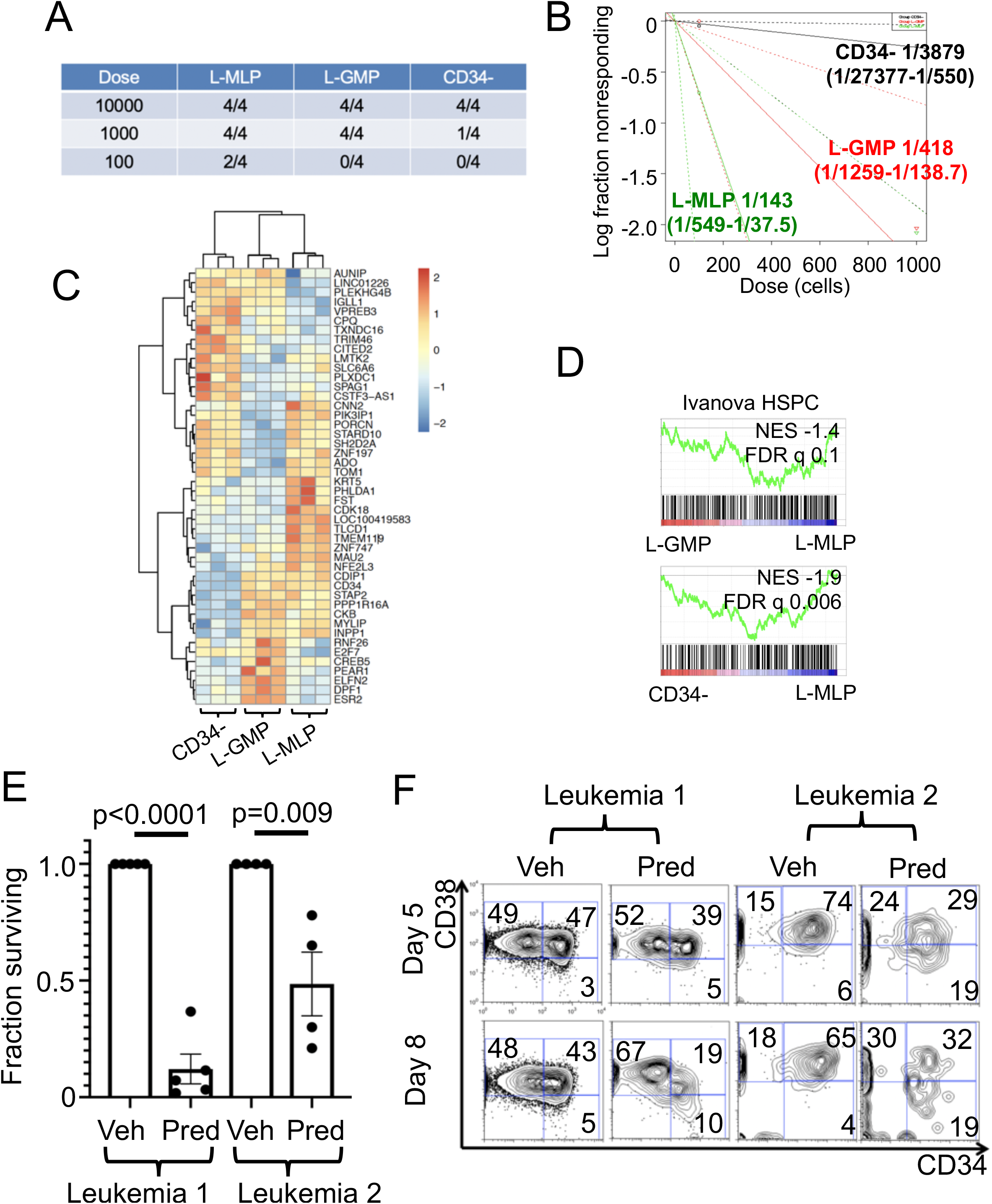
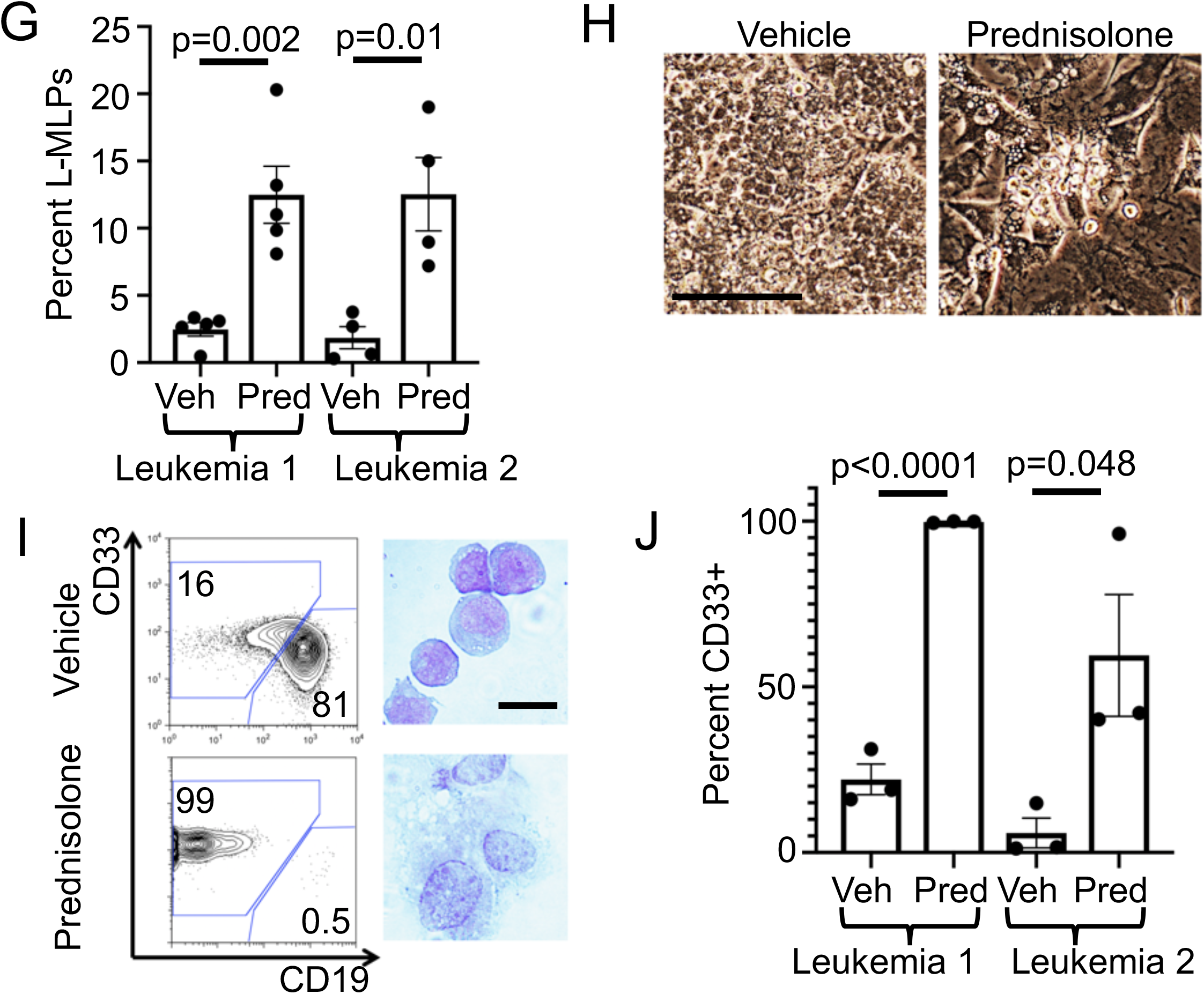

**Figure S4.**
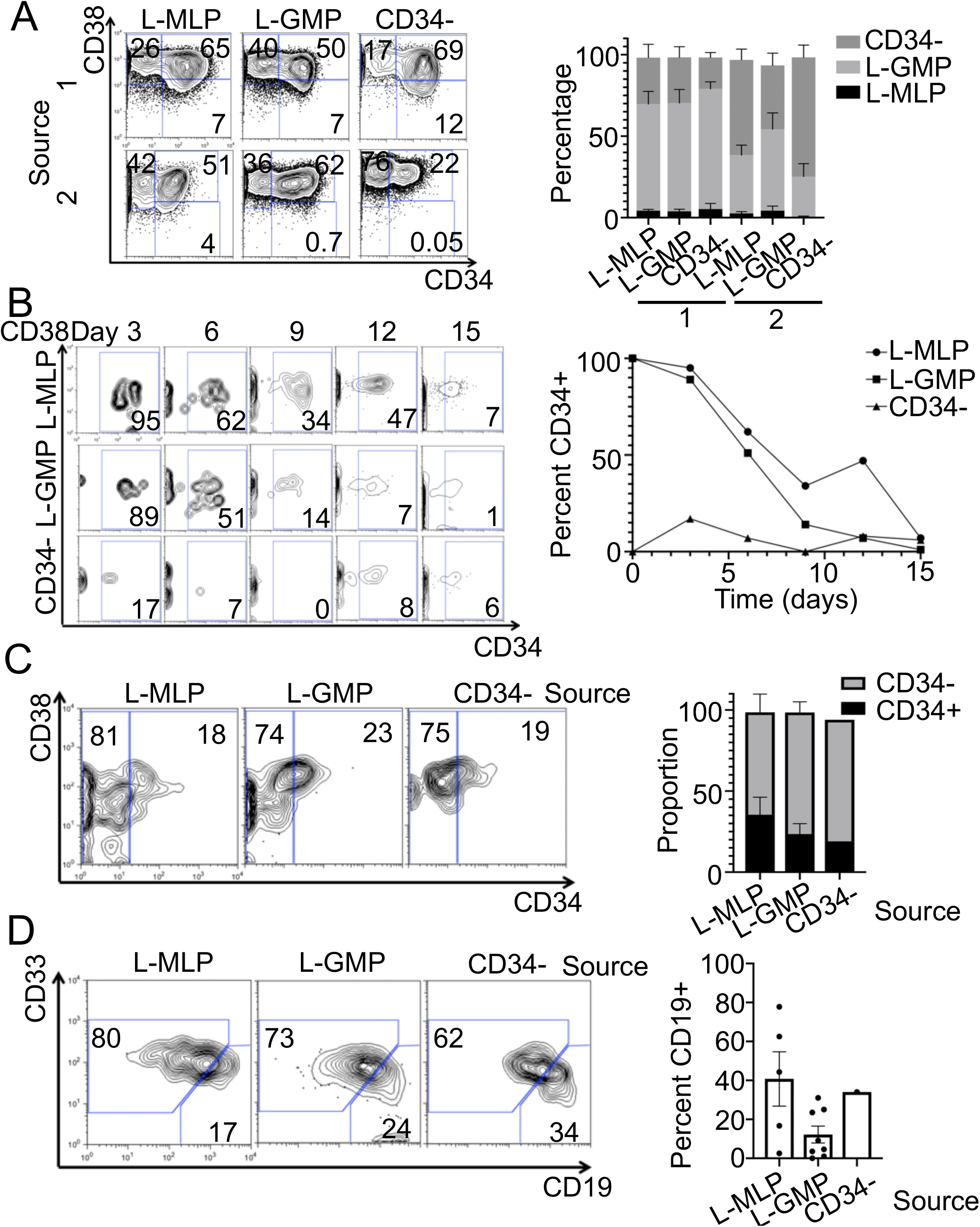

## Notes

### Competing Interest Statement

The authors have declared no competing interest.

### Summary of Updates

Data availability.

