## Supplemental data for "Single-cell analysis uncovers mechanisms of plasticity in leukemia initiating cells"

**SUPPLEMENTAL INFORMATION**

**Fig. S1. Lineage plasticity and scRNA-seq of *MLL*-r B-ALL.**

(A) *MLL*-r B-ALL cells were isolated from leukemic NSG bone marrow and plated on MS5 stroma in the presence of cytokines for four weeks, at which time cultures were analyzed by flow cytometry (gated on human CD45^+^), with CD19^-/low^ CD33^+^ and CD19^+^ CD33^-^ populations isolated by fluorescence activated cells sorting (FACS) and examined by light microscopy following May-Grunwald-Giemsa staining (scale = 10 μm).

(B-C) Single *MLL*-r B-ALL cells were sorted into wells with MS5 stroma and cytokines. After four weeks, clones derived from single cells were picked and analyzed by flow cytometry (gated on human CD45^+^) and the fraction of CD33^+^ and CD19^+^ cells quantified compared to the input leukemia. Results presented as mean ± SEM.

(D) Human CD45+ *MLL*-r B-ALL cells were isolated from a leukemic xenotransplanted mouse and analyzed by single cell RNA sequencing (results shown are for leukemia-2*/MLL-ENL).* t-distributed stochastic neighbor embedding (t-SNE) was used to visualize nine distinct subpopulations.

(E) Heatmap showing gene expression in each of the tSNE clusters.

(F) SingleCellNet was used to classify individual cells relative to normal HSPC and differentiated benchmarks.

(G) The StemID algorithm was used to annotate each single cell with a multipotency score, with results overlaid on the t-SNE plot.

(H) The expression of the indicated cell surface markers in each subpopulation is shown.

(I) Representative results of fluorescence activated cell sorting and purity analysis of each of the assayed leukemia populations.

**Fig. S2. Cellular hierarchy in *MLL*-r B-ALL**.

(A) The indicated cell populations were isolated by FACS from leukemia 2/*MLL-ENL* and cell cycle status analyzed by costaining for Ki67 and DAPI. The proportion of cells in each cell cycle phase is quantified (results are aggregated over 4 independent experiments and nine transplant recipients, presented as mean ± SEM and compared by student’s t-test to L-MLPs, * p <0.05 compared to L-MLP).

(B) The indicated FACS-sorted cell populations were cultured on MS5 stromal layers for 14 days at which time morphology of the cultures was analyzed by phase contrast microscopy (scale = 100 μm).

(C-D) Human/mouse chimerism analysis of leukemia 1/*MLL-AF4* used in Figure 2 xenotransplantation experiments. Results are from mice transplanted with a dose of 400 cells from each population and analyzed either at the onset of terminal leukemia or at the experimental endpoint. Selected flow cytometry plots are presented. Results are aggregated over two independent transplant experiments, presented as mean ± SEM, p values shown, analyzed by unpaired student’s t-test.

(E) In vivo LDA with leukemia 2*/MLL-ENL* was performed with the dose and fraction of transplanted mice developing terminal leukemia presented.

(F) Kaplan-Meier analysis of survival of leukemia 2*/MLL-ENL.* By log-rank test at 100 cell dose, p = 0.04 for L-MLP versus CD34- and p = 0.07 for L-MLP versus L-GMP.

(G) LIC content of each population was quantified by in vivo limiting dilution analysis (see table in (f)). Results are aggregated over two independent xenotransplantation experiments (see Table S2).

(H) Leukemia 2 was engrafted into NSG mice. At the onset of terminal leukemia or at the experimental endpoint (day 150), recipient mice were euthanized and human chimerism quantified (results are presented as mean ± SEM, p = NS for all comparisons by unpaired student’s t-test).

**Fig. S3. Transcriptional and functional heterogeneity in *MLL*-r B-ALL.**

(A-B) The indicated doses of leukemic cells from the peripheral blood of the patient corresponding to xenograft 1 used in this study were transplanted into unconditioned NSG mice, and the incidence of terminal leukemia measured and compared by limiting dilution analysis (for L-MLP versus L-GMP Χ^2^ = 1.29, p = 0.3; for L-MLP versus CD34^-^ Χ^2^ = 10.7, p = 0.001; for L-GMP versus CD34^-^ Χ^2^ = 4.64, p = 0.03).

(C) The indicated cells were sorted and transcript profiles measured by RNA-seq. Heatmap is presented showing gene expression signatures.

(D) Gene set enrichment analysis was used to analyze enrichment for a published hematopoietic stem and progenitor cell (HSPC) signature among the three populations evaluated.

(E) *MLL*-r B-ALL cells were exposed to either 50 μg/ml prednisolone (Pred) or vehicle (Veh) for five days followed by a washout period. Viability after the five-day prednisolone pulse is presented (results compiled five independent experiments for leukemia 1/*MLL-AF4* and four independent experiments for leukemia 2/*MLL-ENL* and presented as mean ± SEM and compared by student’s t-test, p-value shown).

(F) The content of the cultures was periodically analyzed by flow cytometry at the indicated time points with the indicated antibodies.

(G) Content of CD34^+^ CD38^-^ L-MLPs at day 8 following prednisolone or vehicle exposure is shown (results compiled from five independent experiments for leukemia 1 and four independent experiments for leukemia 2 and presented as mean ± SEM and compared by student’s t-test, p-value shown).

(H) Morphology of cells growing on MS5 stromal layers 14 days after prednisolone or vehicle exposure is shown under phase contrast microscopy (scale = 100 μm).

(I) Representative flow cytometry results (left) and morphologic analysis (right, scale = 10 μm) demonstrating a B-lymphoid to myeloid lineage switch in leukemia 1 three weeks following initial prednisolone exposure.

(J) Quantification of CD33^+^ content in three-week cultures from (d) (n = 3 independent experiments analyzed, results presented as mean ± SEM and analyzed by student’s t-test with p value shown).

**Figure S4. Plasticity in leukemia from primary patient cells**.

(A) The indicated leukemic populations were isolated from the peripheral blood of leukemia patients and transplanted into NSG mice. At the onset of morbidity, recipients were euthanized, and the phenotype of the resulting leukemia analyzed by flow cytometry and quantified. Each population from each source was compared by unpaired student’s t-test, with the only significant comparison being leukemia 2 L-GMP versus CD34^-^ source downstream CD34^-^ population where p = 0.01. Results presented at mean± standard error of mean.

(B) The indicated cell populations were sorted from leukemic peripheral blood (corresponding to leukemia 1/*MLL-AF4*) and plated on MS5 stroma with cytokines, and the relative proportion of CD34^+^ cells in the culture was quantified over time. Results are representative of two independent experiments performed with this leukemic specimen.

(C-D) Colonies derived from single cells of the indicated populations grown on MS5 in vitro were analyzed for expression of CD34 (e-f) or CD19 and CD33 (g-h), with the proportions of cells in each gate quantified.
