## Supplementary material for "Single-cell analysis uncovers mechanisms of plasticity in leukemia initiating cells": Table S1

| | Age (years) | Translocation | Rapid heme Panel* | Sex |
|---|---|---|---|---|
| 1 | 0.88 | MLL-AF4 | KRAS c.35G>A p.G12D 50.9% VAF | F |
| 2 | 0.27 | MLL-ENL | KRAS c.35G>A 42.5% VAF; CNV+X | M |
| 3 | 16.09 | MLL-EPS15 | PDS5B c.4123C>T p.R1375* 23.9% VAF; CNV +X | M |

*Technique developed in Kluk et al., 2016
