## Supplementary material for "Single-cell analysis uncovers mechanisms of plasticity in leukemia initiating cells": Table S2 part 1

Table S2
Leukemia 1, MLL-AF4 patient derived xenograft in vivo xenotransplants

Experiment 1, engrafted 1/19/2018

| Source | Dose | Number | DOD | Days |
|---|---|---|---|---|
| L-GMP | 4000 | 1 | 3/22/18 | 61 |
| L-GMP | 4000 | 2 | 3/25/18 | 64 |
| L-GMP | 4000 | 3 | 3/25/18 | 64 |
| L-GMP | 4000 | 4 | 3/25/18 | 64 |
| L-MLP | 4000 | 1 | 4/9/18 | 79 |
| L-MLP | 4000 | 2 | 4/9/18 | 79 |
| L-MLP | 4000 | 3 | 4/9/18 | 79 |
| L-MLP | 4000 | 4 | 4/9/18 | 79 |
| CD34- | 4000 | 1 | 4/17/18 | 87 |
| CD34- | 4000 | 2 | 17-Apr | 87 |
| CD34- | 4000 | 3 | 4/17/18 | 87 |
| CD34- | 4000 | 4 | 4/17/18 | 87 |
| L-GMP | 40000 | 1 | 3/26/18 | 65 |
| L-GMP | 40000 | 2 | 3/26/18 | 65 |
| L-GMP | 400 | 1 | 4/18/18 | 88 |
| L-GMP | 400 | 2 | Survived | |
| L-GMP | 400 | 3 | Survived | |
| L-GMP | 400 | 4 | Survived | |
| L-MLP | 400 | 1 | 4/18/18 | 88 |
| L-MLP | 400 | 2 | 4/21/18 | 91 |
| L-MLP | 400 | 3 | 4/23/18 | 93 |
| L-MLP | 400 | 4 | 4/23/18 | 93 |
| CD34- | 400 | 1 | 4/26/18 | 96 |
| CD34- | 400 | 2 | Survived | |
| CD34- | 400 | 3 | Survived | |
| CD34- | 400 | 4 | Survived | |
| CD34- | 40000 | 1 | 3/25/18 | 64 |

Experiment 2, engrafted 2/13/2019

| Source | Dose | Number | DOD | Days |
|---|---|---|---|---|
| L-MLP | 40 | 1 | 5/30/19 | 101 |
| L-MLP | 40 | 2 | 6/2/19 | 103 |
| L-MLP | 40 | 3 | 6/2/19 | 106 |
| L-MLP | 40 | 4 | Survived | |
| L-MLP | 400 | 1 | 6/21/19 | 123 |
| L-MLP | 400 | 2 | 6/22/19 | 124 |
| L-MLP | 400 | 3 | 6/22/19 | 124 |
| L-MLP | 400 | 4 | 5/30/19 | 101 |
| L-GMP | 40 | 1 | 6/21/19 | 123 |
| L-GMP | 40 | 2 | Survived | |
| L-GMP | 40 | 3 | Survived | |
| L-GMP | 40 | 4 | Survived | |
| CD34- | 40 | 1 | 7/3/19 | 135 |
| CD34- | 40 | 2 | Survived | |
| CD34- | 400 | 1 | 6/21/19 | 123 |
| CD34- | 400 | 2 | 6/22/19 | 124 |
| CD34- | 400 | 3 | 6/29/19 | 131 |
| CD34- | 400 | 4 | 7/6/19 | 138 |
| L-GMP | 400 | 1 | 6/22/19 | 124 |
| L-GMP | 400 | 2 | 6/22/19 | 124 |
| L-GMP | 400 | 3 | 6/22/19 | 124 |
| L-GMP | 400 | 4 | 6/23/19 | 125 |

Aggregate over two experiments:
for L-MLP versus L-GMP $X^2$ = 10.7, p = 0.001
for L-MLP versus CD34- X2 = 10.2, p = 0.001
for L-GMP versus CD34- X2 = 0.005, p = 0.9
