## Supplementary material for "Single-cell analysis uncovers mechanisms of plasticity in leukemia initiating cells": Table S2 part 2

Leukemia 2, MLL-ENL patient derived xenograft in vivo xenotransplants

Experiment 1, engrafted 2/27/2019

| Source | Dose | Number | DOD | Days |
|---|---|---|---|---|
| CD34- | 1000 | 1 | 6/13/19 | 106 |
| CD34- | 1000 | 2 | 6/17/19 | 110 |
| CD34- | 1000 | 3 | 6/17/19 | 110 |
| CD34- | 1000 | 4 | 6/18/19 | 111 |
| L-MLP | 1000 | 1 | 6/17/19 | 110 |
| L-MLP | 1000 | 2 | 6/17/19 | 110 |
| L-MLP | 1000 | 3 | 6/18/19 | 111 |
| L-MLP | 1000 | 4 | 6/18/19 | 111 |
| L-GMP | 1000 | 1 | 7/6/19 | 129 |
| L-GMP | 1000 | 2 | 7/14/19 | 137 |
| L-GMP | 1000 | 3 | 7/15/19 | 138 |
| L-GMP | 1000 | 4 | 7/15/19 | 138 |
| CD34- | 100 | 1 | survived | |
| CD34- | 100 | 2 | survived | |
| CD34- | 100 | 3 | survived | |
| CD34- | 100 | 4 | survived | |
| L-MLP | 100 | 1 | 7/4/19 | 127 |
| L-MLP | 100 | 2 | 7/21/19 | 144 |
| L-MLP | 100 | 3 | 7/23/19 | 146 |
| L-MLP | 100 | 4 | survived | |
| L-GMP | 100 | 1 | survived | |
| L-GMP | 100 | 2 | survived | |
| L-GMP | 100 | 3 | survived | |
| L-GMP | 100 | died shortly after transplant - not leukemic death and so excluded | | |

Experiment 2, engrafted 5/14/2019

| Source | Dose | Number | DOD | Days |
|---|---|---|---|---|
| CD34- | 4000 | 1 | 8/9/19 | 87 |
| CD34- | 4000 | 2 | 8/16/19 | 94 |
| CD34- | 4000 | 3 | 8/17/19 | 95 |
| CD34- | 4000 | 4 | 8/25/19 | 103 |
| L-MLP | 4000 | 1 | 8/9/19 | 87 |
| L-MLP | 4000 | 2 | 8/16/19 | 94 |
| L-MLP | 4000 | 3 | 8/19/19 | 97 |
| L-MLP | 4000 | 4 | 8/20/19 | 98 |
| L-GMP | 4000 | 1 | 8/19/19 | 97 |
| L-GMP | 4000 | 2 | 8/27/19 | 105 |
| L-GMP | 4000 | 3 | 8/27/19 | 105 |
| L-GMP | 4000 | 4 | 8/30/19 | 108 |
| CD34- | 400 | 1 | 8/19/19 | 97 |
| CD34- | 400 | 2 | 9/24/19 | 133 |
| CD34- | 400 | 3 | survived | |
| CD34- | 400 | 4 | survived | |
| L-MLP | 400 | 1 | 9/2/19 | 111 |
| L-MLP | 400 | 2 | 9/4/19 | 113 |
| L-MLP | 400 | 3 | survived | |
| L-MLP | 400 | 4 | survived | |
| L-GMP | 400 | 1 | 9/17/19 | 126 |
| L-GMP | 400 | 2 | survived | |
| L-GMP | 400 | 3 | survived | |
| L-GMP | 400 | 4 | survived | |

Aggregate over two experiments:
for L-MLP versus L-GMP $X^2 = 2.2$, $p = 0.1$
for L-MLP versus CD34- X2 = 1.1, p = 0.3;
for L-GMP versus CD34- X2 = 0.2, p = 0.7
